# Enhancing the Efficacy of Oncolytic Viruses by Engineering Virus–Host Interactions Using CRISPR-Cas13d

**DOI:** 10.64898/2026.11.20.768134

**Authors:** Zahra Ameli, Nika Kooshki Zamani, Zumama Khalid, Reza Rezaei, Gabriel St-Laurent, Leila Khani Khouzani, Rémi Cormier, Phuong Anh Nguyen, Pedram Abdollahpour, Maria Fernanda Solorza Reyes, Georgia Cheung, Shervin Taslimi, Kian Jafari, Taha Azad

**Author notes:** **Corresponding Author** *Taha Azad,* Department of Microbiology and Infectious Diseases, Faculty of Medicine and Health Sciences, Université de Sherbrooke, Sherbrooke, Québec, Canada.

## Abstract

Oncolytic herpes simplex virus type 1 (oHSV-1) is a promising cancer therapy, but tumor-cell antiviral responses can restrict viral replication. Strategies for temporally controlled targeting of antiviral factors remain limited. Here, we developed an inducible CRISPR-Cas13d platform to modulate host responses during HSV-1 infection. We screened 13 pharmacological inhibitors across tumor models to identify pathways associated with restricted infection. Reporter assays identified Cas13d as the preferred effector over Cas13b, while endogenous-target experiments demonstrated Cas13d-mediated reductions in target protein abundance. We optimized guide RNA expression using RNA polymerase III promoters, selected p7SK, and established doxycycline-responsive Tet-On and Tet-Off systems. The platform was incorporated into HSV-1 to generate a gRNA library targeting 15 host antiviral genes, together with two non-targeting controls. Doxycycline-dependent reporter readouts demonstrated inducible activity. Evaluation in EMT6, CT26, and 786-O cells revealed guide-dependent differences in fluorescence. These findings establish a programmable framework for investigating virus–host interactions during oncolytic HSV infection.

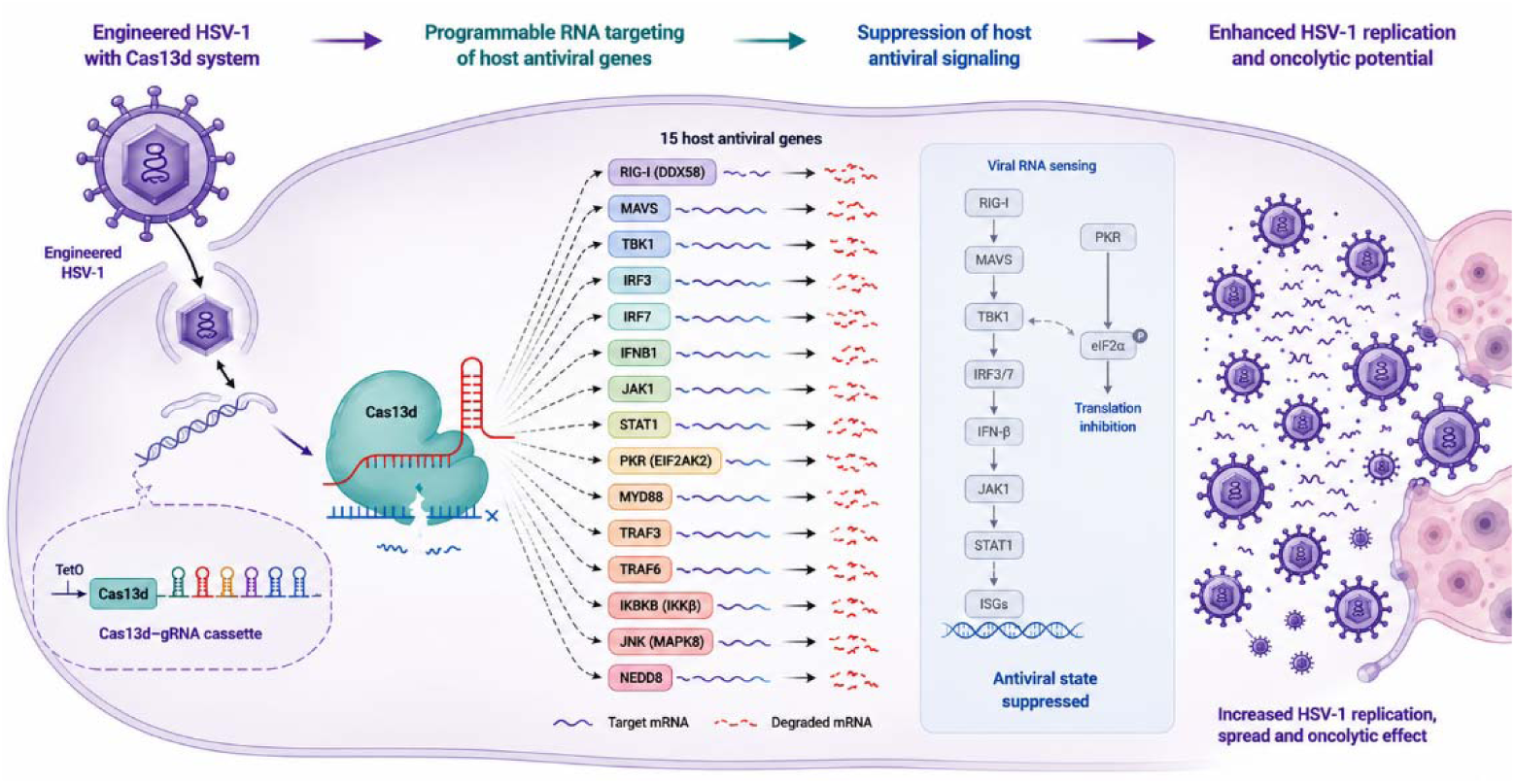

## INTRODUCTION

Cancer remains a major global health challenge despite substantial advances in surgery, radiotherapy, chemotherapy, targeted therapy, and immunotherapy. Tumor heterogeneity, treatment resistance, and immune evasion continue to limit durable responses in many malignancies, creating a need for approaches that can eliminate cancer cells while promoting effective antitumor immunity ^1–3^.

Oncolytic virotherapy addresses this need by using naturally occurring or genetically engineered viruses to preferentially infect and destroy tumor cells while stimulating antitumor immune responses. In addition to direct oncolysis, some oncolytic viruses can induce immunogenic cell death, accompanied by the exposure or release of damage-associated molecular patterns (DAMPs), including calreticulin, ATP, and HMGB1, which support antigen uptake and presentation ^4,5^. These properties provide a rationale for combining virotherapy with other immunotherapeutic approaches ^6–9^. However, antiviral defenses in infected tumor cells can restrict viral replication and intratumoral spread, while innate responses also contribute to the immune effects of treatment ^10,11^.

Herpes simplex virus type 1 (HSV-1) is a widely investigated oncolytic platform because of its broad cellular tropism, large genome, well-characterized biology, and amenability to genetic modification^12,13^. The clinical development of talimogene laherparepvec (T-VEC), an engineered HSV-1 expressing granulocyte-macrophage colony-stimulating factor (GM-CSF), established the therapeutic potential of this platform ^14,15^. Nevertheless, responses to oHSV vary among tumor types, reflecting differences in tumor-cell susceptibility and intrinsic antiviral signaling ^16^. These differences suggest that the host factors restricting viral replication may not be identical across tumor contexts. Outcomes are also shaped by the immune state and molecular characteristics of the infected tumor, complicating the prediction of therapeutic responses.

HSV infection is detected by cellular innate immune sensors that activate interconnected antiviral pathways. Viral DNA sensing through factors such as IFI16 and the cGAS– STING pathway can promote TBK1-dependent activation of interferon regulatory factors and type I interferon production ^17–19^. Type I interferons signal through IFNAR and the JAK/STAT pathway to induce interferon-stimulated genes, establishing an antiviral state ^20^. Among the downstream effectors, protein kinase R (PKR; EIF2AK2) restricts viral protein synthesis through phosphorylation of eIF2α ^21^. Additional regulators, including RIG-I/MAVS and NF-κB-associated signaling, further shape the response. Because these pathways are interconnected and their activity differs between cell types, the contribution of individual host factors to oHSV restriction is context dependent.

Several approaches have been explored to overcome these antiviral barriers. Jackson et al. (2016) showed that pharmacological inhibition of STAT1-and NF-κB-associated signaling reduced basal interferon-stimulated gene expression and improved productive oHSV infection in malignant peripheral nerve sheath tumor cells ^22^. In a complementary virus-engineering strategy, Hong et al. reported that an oHSV expressing a PKR-targeting short hairpin RNA altered antiviral signaling and improved oncolytic immune therapy in preclinical models ^23^. These studies support host-directed modulation as a means of influencing viral activity. However, small-molecule inhibitors can affect multiple cellular processes, while targeting a single factor may not address the diversity of antiviral mechanisms across tumors. A programmable strategy that can be redirected toward different host transcripts would provide a flexible framework for investigating and modulating these interactions.

CRISPR-Cas13 systems offer programmable RNA targeting without directly altering genomic DNA ^24^. Cas13d is a compact RNA-guided ribonuclease that can be adapted for transcriptome engineering in mammalian cells ^25^. Its potential for biomedical and synthetic biology applications has been reviewed by Palaz et al. (2021), Yang and Patel (2024), and Hou et al. (2026) ^26–28^. However, Cas13-mediated silencing depends on guide sequence, target accessibility, and effector abundance. Large-scale screens have shown that guide performance varies substantially and that both on-target and off-target activity should be considered during guide selection ^29^. Recent work has further highlighted the importance of controlling Cas13d expression to manage collateral RNA cleavage ^30^. These considerations make optimization of both guide design and effector expression important for a reliable viral platform.

Temporal regulation is particularly relevant when host antiviral genes are targeted during oncolytic infection. Although transient suppression may support viral amplification, sustained interference with innate pathways could also affect immune functions that contribute to tumor control. Tetracycline-responsive Tet-On and Tet-Off systems provide established methods for regulating transgene expression in response to doxycycline ^31,32^. Incorporating such control into an HSV-encoded Cas13d system could allow the timing of effector expression to be adjusted during infection.

In this study, we developed an inducible CRISPR-Cas13d platform to engineer virus–host interactions in HSV-1. We first evaluated host antiviral pathways through pharmacological screening across tumor cell models. We then compared Cas13b and Cas13d activity, assessed endogenous RNA targeting, optimized guide RNA expression, and established doxycycline-regulated control of Cas13d. Finally, we generated a gRNA library targeting 15 host antiviral genes, along with two non-targeting controls, and evaluated the resulting HSV-iCas13d constructs in tumor cell models.

We hypothesized that inducible Cas13d-mediated targeting would provide a programmable approach to modulating host antiviral responses and improving the replicative activity of oncolytic HSV.

## RESULTS AND DISCUSSION

To identify host pathways associated with restricted HSV replication and establish a basis for their targeted modulation, we combined transcriptomic profiling and pharmacological screening with CRISPR-Cas13d-based viral engineering. This approach integrates chemical modulation of host signaling pathways using small-molecule inhibitors with molecular and synthetic biology strategies to develop a programmable system for investigating and engineering virus–host interactions.

### Transcriptomic profiling reveals distinct host responses to oncolytic HSV infection across cancer cell lines

To characterize host transcriptional responses to HSV infection and identify pathways potentially involved in antiviral defense, we performed RNA sequencing (RNA-seq) in four murine cancer cell lines: EMT6, CT26, AT3, and 67NR. Each cell line was evaluated under two experimental conditions: uninfected control and oncolytic HSV-1 (oHSV-1) infection at a multiplicity of infection (MOI) of 1. Following infection, cells were incubated for 72 h at 37 °C under 5% CO before collection and RNA stabilization. The resulting samples were subjected to RNA sequencing and subsequent differential gene expression and pathway enrichment analyses **(Figure 1a)**.

**Figure 1.**
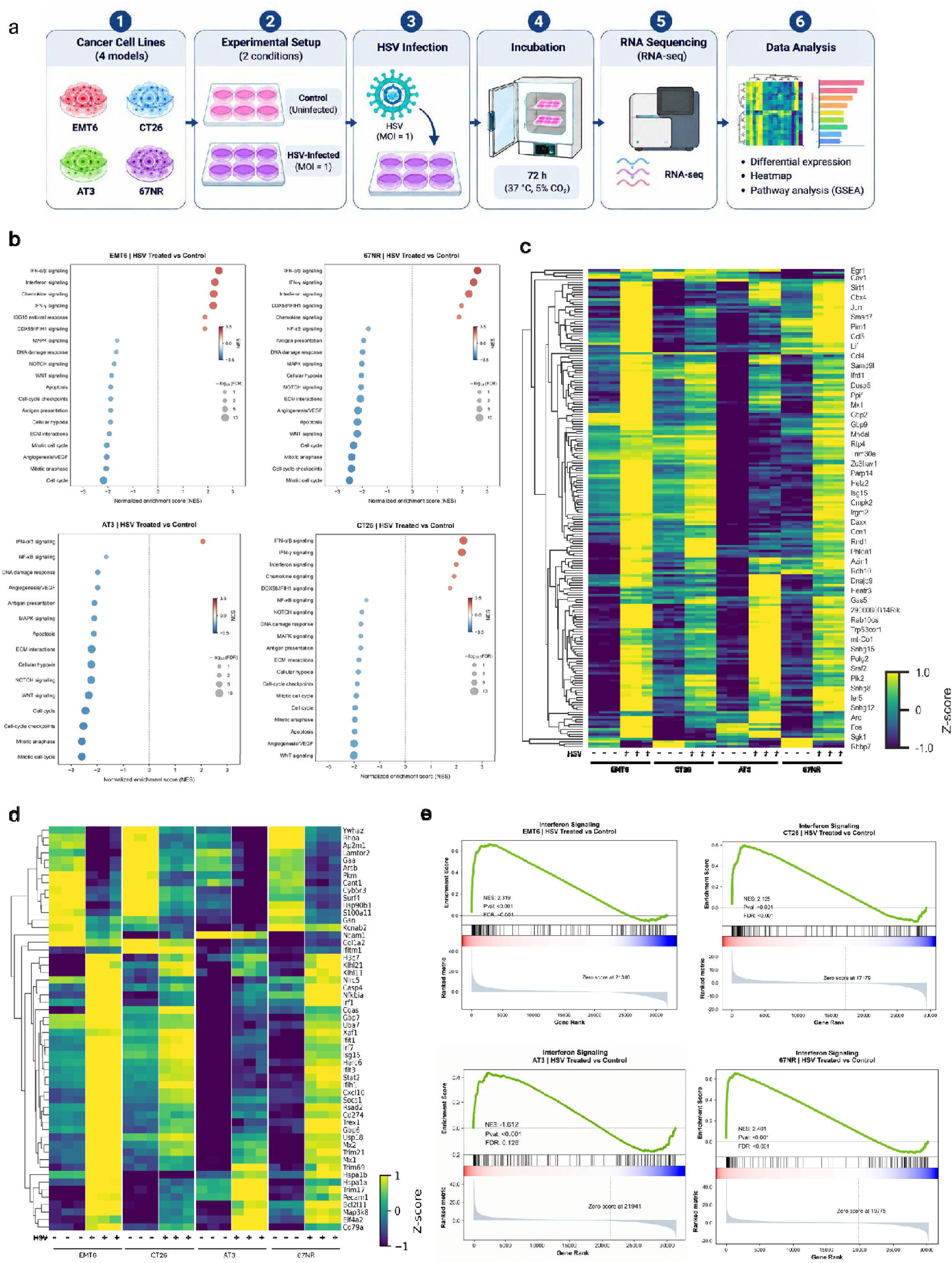
Transcriptomic analysis of cancer cell responses to oHSV-1 infection. (a) Schematic representation of the experimental workflow using EMT6, CT26, AT3, and 67NR cells infected with oHSV-1 at an MOI of 1, followed by 72 h incubation and RNA sequencing. (b) GSEA dot plots showing enriched biological pathways in each cell line. (c, d) Heatmaps displaying global gene expression patterns and selected immune-related genes, respectively, across control and HSV-infected samples. (e) GSEA enrichment plots showing interferon signaling in the four cell lines. Gene expression in the heatmaps is presented as row-wise Z-scores. NES, normalized enrichment score; FDR, false discovery rate.

Gene set enrichment analysis (GSEA) revealed substantial transcriptional changes across several biological pathways associated with innate immunity and antiviral responses ^17^**(Figure 1b)**. Interferon-related pathways, including IFN-α signaling, IFN-γ signaling, and interferon signaling, showed positive enrichment across multiple cell lines. Chemokine signaling and other immune-associated pathways were also positively enriched^33^. In contrast, several pathways associated with cellular proliferation, including cell-cycle regulation, mitotic cell cycle, and cell-cycle checkpoints, displayed negative enrichment scores. Additional changes were observed in pathways related to NF-κB signaling, antigen presentation, MAPK signaling, DNA damage responses, NOTCH signaling, and extracellular matrix interactions. Importantly, the enrichment patterns varied among the four cell lines, indicating that the transcriptional response to HSV infection was influenced by the cellular background.

To further characterize these changes at the individual gene level, we examined hierarchical clustering heatmaps displaying global differentially expressed genes and selected immune-associated genes across control and HSV-infected samples **(Figure 1c,d)**. The global expression heatmap revealed distinct transcriptional profiles between the two experimental conditions, with considerable variation among the four cell lines **(Figure 1c)**. Several genes exhibited increased expression following HSV infection, whereas others showed reduced expression. Examination of selected immune-associated genes further revealed changes in multiple genes involved in innate immune recognition, inflammatory signaling, and antiviral responses **(Figure 1d)**. These patterns highlighted the heterogeneous nature of host immune responses and provided a more focused view of the genes contributing to the pathway-level changes observed following infection. Representative genes included interferon-associated factors such as Ifit1, Ifit3, Isg15, Mx1, Mx2, Irf1, Irf7, and Stat2 ^34^.

Given the prominent enrichment of interferon-associated pathways, we next assessed the coordinated behavior of interferon signaling using GSEA^35^ in each cell line **(Figure 1e)**. Positive normalized enrichment scores (NES) were observed in EMT6 (NES = 2.319), CT26 (NES = 2.125), AT3 (NES = 1.612), and 67NR (NES = 2.401). The enrichment was statistically significant in EMT6, CT26, and 67NR (FDR < 0.001), whereas the enrichment in AT3 did not reach statistical significance (FDR = 0.126). Thus, although interferon-associated transcriptional responses were observed across all four models, statistically significant pathway-level enrichment was detected in only three. These findings indicate that the magnitude and coordination of interferon-associated responses differed among the cancer cell lines.

Additional transcriptomic analyses were performed to further characterize changes in host cellular programs following HSV infection (Supplementary Figure S1). GSEA revealed negative enrichment scores for apoptosis-related gene sets in EMT6 (NES = −1.987), CT26 (NES = −1.941), AT3 (NES = −2.006), 67NR (NES = −2.026), and the human 786-O cell line (NES = −1.754) **(Supplementary Figure S1a)**. All five enrichments were statistically significant, with FDR values below 0.001 for EMT6, CT26, AT3, and 67NR, and an FDR of 0.012 for 786-O. Given the HSV-treated-versus-control comparison, these negative enrichment scores indicate that apoptosis-associated genes were preferentially distributed toward the negative end of the ranked gene lists, consistent with relatively lower expression following HSV infection. However, these transcriptomic findings alone do not establish functional changes in apoptosis.

Furthermore, additional heatmap analyses revealed distinct expression patterns among selected genes associated with innate immune responses, interferon signaling, cytokine signaling, and antiviral pathways across the analyzed samples **(Supplementary Figure S1b)**. These heatmaps provided complementary information on the expression variability of individual immune-associated genes and further illustrated the heterogeneous transcriptional responses associated with HSV infection.

These observations highlight that HSV-1 infection induces distinct, cell-dependent transcriptional responses, particularly involving interferon-associated and other immune-related pathways. The observed differences in pathway enrichment and gene expression highlight the variability of intrinsic antiviral responses across tumor cell models. This transcriptomic characterization provided a basis for the subsequent pharmacological investigation of host antiviral pathways and the identification of candidate factors for targeted modulation using the engineered CRISPR-Cas13d platform.

### Pharmacological targeting of host antiviral signaling enhances HSV replication across tumor cell models

Because host antiviral signaling can restrict HSV replication, we first investigated whether pharmacological modulation of distinct antiviral pathways could enhance viral replication in tumor cells ^22,36^. A panel of 13 small-molecule inhibitors targeting different components of host antiviral signaling was evaluated, including MRT67307, GSK8612, momelotinib, C25-140, amlexanox, a PKR inhibitor (PKRi), BX795, dimethyl fumarate, pevonedistat, BAY 11-7082, ruxolitinib, TPCA-1, and 4-octyl itaconate (4-OI) **(Figure 2a)**. Their effects on HSV replication were assessed across five tumor cell lines representing distinct cancer models: EMT6, CT26, 786-O, 67NR, and AT3.

**Figure 2.**
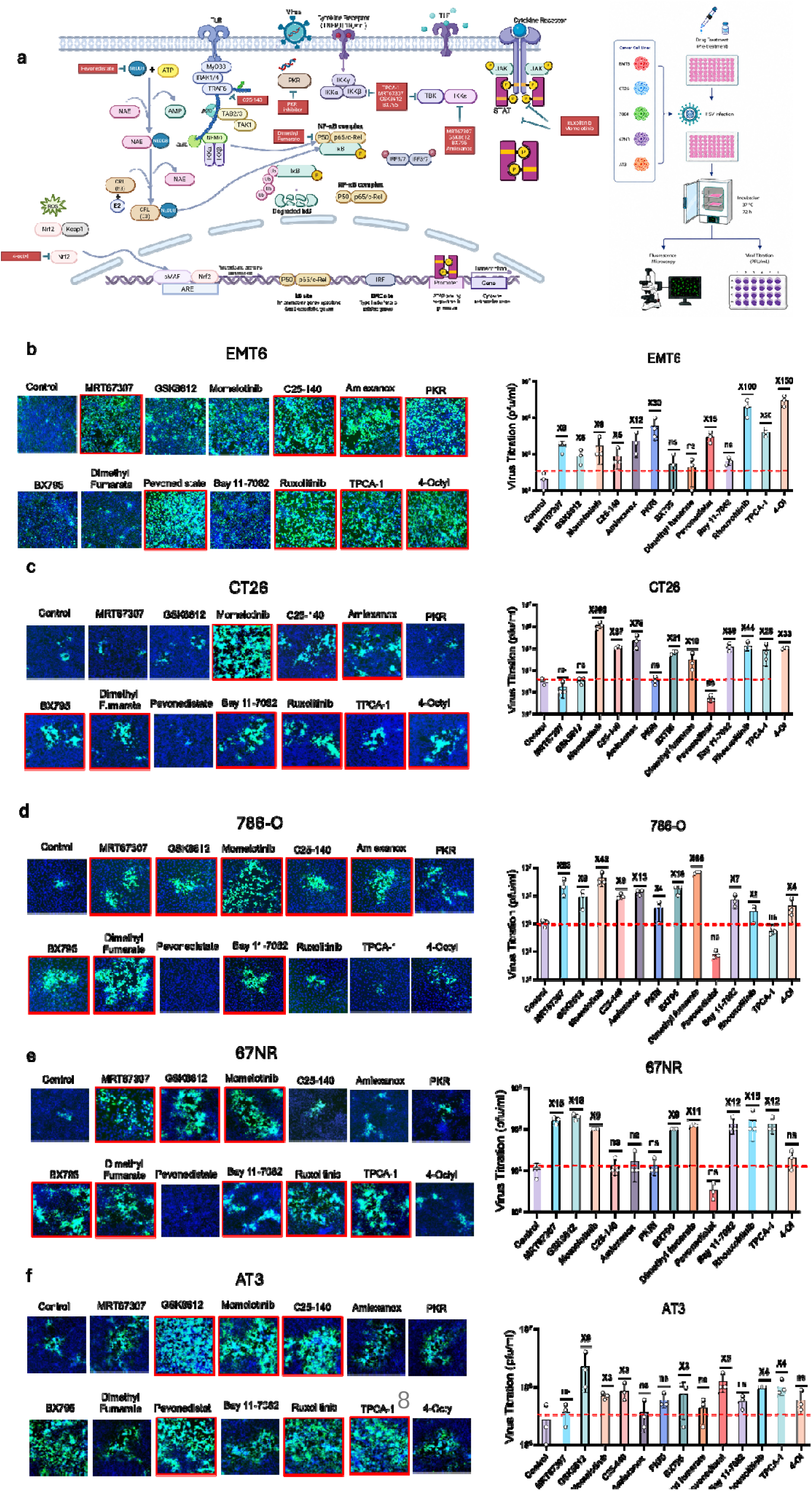
Pharmacological inhibition of host antiviral signaling enhances HSV replication across tumor cell models. (a) Schematic overview of the host antiviral signaling pathways targeted in the pharmacological screen and the experimental workflow. (b–f) Representative fluorescence images and infectious viral titers following treatment of EMT6 (b), CT26 (c), 786-O (d), 67NR (e), and AT3 (f) cells. Cells were pretreated with the indicated inhibitors (0.1 µM) or DMSO vehicle control before infection with recombinant HSV-1 at an MOI of 0.1. At 72 h post-infection, viral reporter fluorescence was assessed by microscopy, and infectious virus production was quantified by plaque assay. Images were acquired under identical imaging conditions. Viral titers are presented as mean ± SD (n = 3 independent biological replicates) in plaque-forming units per milliliter (PFU/mL). Red dashed lines indicate the infectious viral titers in the corresponding control groups. Fold increases relative to the DMSO-treated control are indicated above the corresponding bars. The inhibitors included MRT67307, GSK8612, momelotinib, C25-140, amlexanox, PKR inhibitor (PKRi), BX795, dimethyl fumarate, pevonedistat, BAY 11-7082, ruxolitinib, TPCA-1, and 4-octyl itaconate (4-OI). ns, not significant.

Cells were pretreated with the indicated inhibitors before infection with recombinant HSV-1. At 72 h post-infection, viral reporter fluorescence was evaluated by fluorescence microscopy, and infectious virus production was quantified by plaque assay. In EMT6 cells, several inhibitors enhanced viral replication relative to control treatment, with the strongest effects observed following treatment with 4-OI (∼150-fold) and ruxolitinib (∼100-fold) **(Figure 2b)**. These increases were accompanied by enhanced reporter fluorescence and higher infectious viral titers, indicating that pharmacological modulation of host antiviral signaling can markedly enhance HSV production in this model.

A particularly pronounced response was observed in CT26 cells, where momelotinib increased infectious viral titers by approximately 389-fold, representing the largest enhancement observed across the five tumor models. C25-140 produced the second strongest response (∼78-fold) **(Figure 2c)**. These findings suggest that CT26 cells are particularly responsive to pharmacological disruption of host antiviral signaling, with JAK/STAT pathway inhibition producing the most pronounced effect in this model.

In 786-O cells, dimethyl fumarate produced the strongest enhancement in viral titer (∼65-fold), followed by momelotinib (∼42-fold) **(Figure 2d)**. This response differed from those observed in EMT6 and CT26 cells, where 4-OI/ruxolitinib and momelotinib, respectively, produced the strongest effects. These findings further highlight the cell-dependent contribution of distinct host signaling pathways to HSV restriction.

In 67NR cells, the overall magnitude of enhancement was more moderate. Nevertheless, several inhibitors increased viral replication relative to control treatment **(Figure 2e)**. GSK8612 produced the strongest increase in viral titer (∼18-fold), followed by MRT67307 and ruxolitinib (∼15-fold each). These results indicate that pharmacological inhibition of antiviral signaling can also enhance HSV replication in 67NR cells, although the magnitude of the response was lower than that observed in CT26 and 786-O cells.

Similarly, in AT3 cells, GSK8612 produced the largest increase in viral titer (∼9-fold), followed by pevonedistat (∼5-fold) **(Figure 2f)**. Compared with the other tumor models, AT3 cells showed a more modest overall response to pharmacological pathway inhibition, further suggesting that the contribution of individual host pathways to antiviral restriction varies across tumor cell types.

The findings suggest that pharmacological modulation of host antiviral signaling can enhance infectious HSV production across multiple tumor cell models, while revealing substantial differences in the magnitude and pathway dependence of these responses. Notably, JAK/STAT pathway inhibition emerged as a recurrent feature of the strongest responses, with ruxolitinib or momelotinib among the most effective treatments in EMT6, CT26, and 67NR cells ^22^. However, other inhibitors, particularly GSK8612 and dimethyl fumarate, produced the strongest responses in specific cellular backgrounds. These findings emphasize the context-dependent nature of host antiviral restriction and support the development of more precise and programmable approaches to modulating individual host antiviral factors using CRISPR-Cas13d. Importantly, the effects of pharmacological inhibitors may involve multiple cellular targets and therefore do not independently establish the contribution of individual host genes ^37^.

### Functional characterization of Cas13 variants and optimization of the CRISPR-Cas13d system

To establish an effective RNA-targeting platform for subsequent incorporation into the oncolytic HSV genome, we first compared the activities of Cas13b and Cas13d using fluorescence-and luminescence-based reporter assays in HEK293 cells. Cells were transfected with Cas13b-or Cas13d-expressing constructs together with a BFP-NanoLuc (BFP-Nluc) reporter system **(Figure 3a)** ^25^. Reporter suppression was evaluated by monitoring BFP fluorescence and NanoLuc luminescence following Cas13-mediated RNA cleavage.

**Figure 3.**
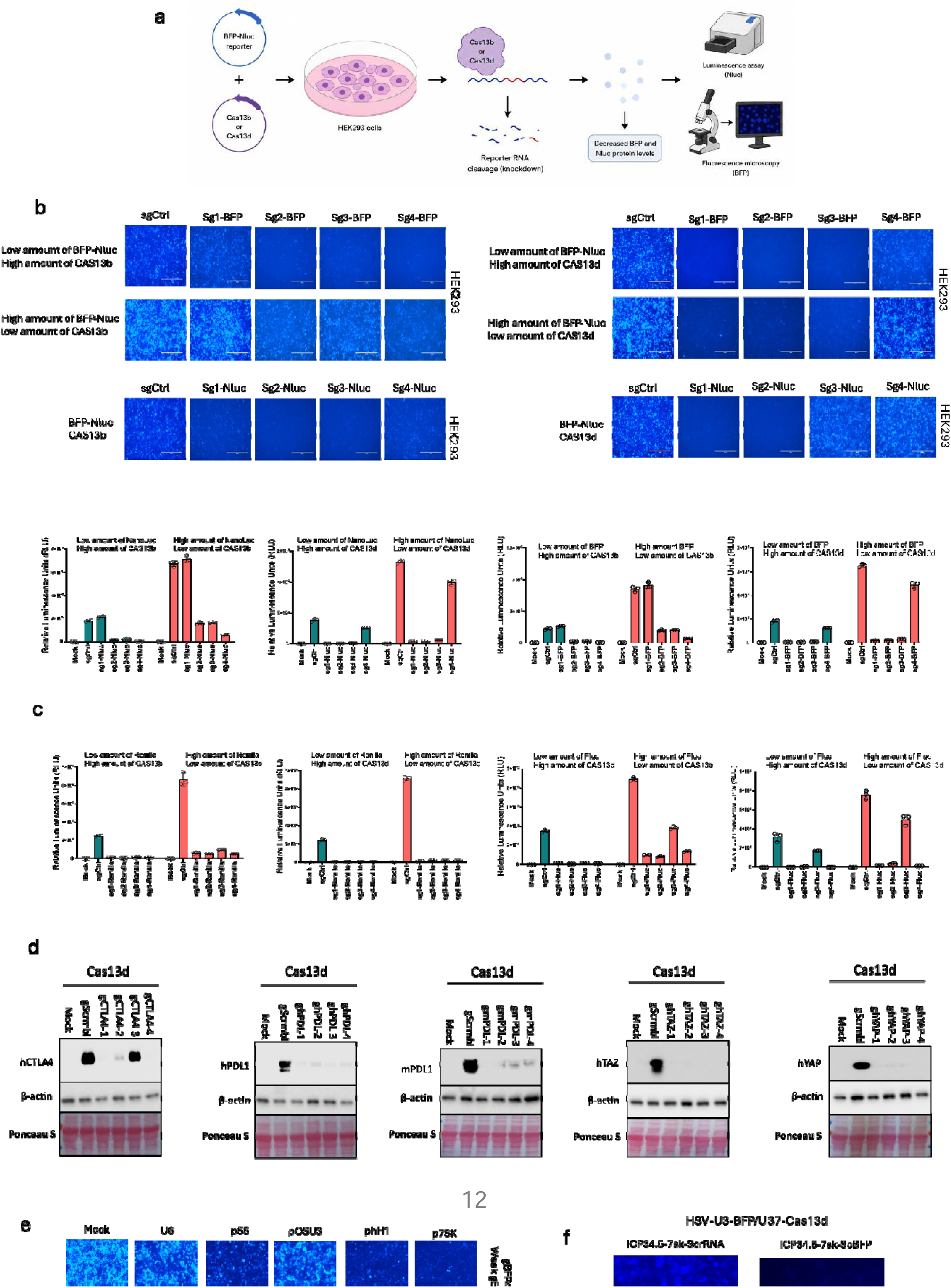
Functional characterization of Cas13b and Cas13d and optimization of the CRISPR-Cas13d system. (a) Schematic representation of the reporter-based strategy used to compare Cas13b and Cas13d activity in HEK293 cells. (b) Representative fluorescence images showing BFP reporter expression and corresponding luminescence measurements of BFP-NanoLuc reporter suppression using four targeting gRNAs (sg1–sg4) and a non-targeting control (sgCtrl) under different reporter and Cas13 expression conditions. (c) Comparison of Cas13b and Cas13d targeting activity using Renilla and firefly luciferase (Fluc) reporters. (d) Western blot analysis of Cas13d-mediated targeting of human CTLA-4 (hCTLA-4), human PD-L1 (hPD-L1), mouse PD-L1 (mPD-L1), human TAZ (hTAZ), and human YAP (hYAP). β-Actin was used as a loading control, and Ponceau S staining was used to assess total protein loading and membrane transfer. (e) Comparison of U6, p5S, pOSU3, pH1, and p7SK promoters for gRNA expression under weak and strong BFP reporter expression conditions. (f) Representative fluorescence images showing BFP reporter suppression in HEK293 cells expressing the HSV-U3-BFP/U37-Cas13d construct containing either the non-targeting control cassette (ICP34.5-7SK-scrRNA) or the BFP-targeting gRNA cassette (ICP34.5-7SK-sgBFP).

We initially assessed the targeting of NanoLuc and BFP reporter sequences using four guide RNAs (sg1–sg4) and a non-targeting control (sgCtrl). Fluorescence microscopy revealed reduced BFP signals with several targeting guides compared with the non-targeting control, although the magnitude of suppression varied among guides and experimental conditions **(Figure 3b)**. The corresponding luminescence measurements also showed guide-dependent differences in reporter activity. For NanoLuc targeting, Cas13b-mediated suppression varied across the four guides. Under the lower-reporter/higher-Cas13b condition, sg2-Nluc, sg3-Nluc, and sg4-Nluc substantially reduced luminescence, whereas sg1-Nluc showed comparatively limited suppression. A similar guide-dependent pattern was observed under the higher-reporter/lower-Cas13b condition. Cas13d also showed guide-dependent activity against NanoLuc. In particular, sg1-Nluc, sg2-Nluc, and sg3-Nluc markedly reduced luminescence under both tested expression conditions, whereas sg4-Nluc retained comparatively higher residual reporter activity.

BFP targeting similarly revealed differences in guide performance. Under the lower-BFP/higher-Cas13b condition, sg2-BFP, sg3-BFP, and sg4-BFP reduced reporter signals relative to sgCtrl, whereas sg1-BFP showed limited suppression. Under the higher-BFP/lower-Cas13b condition, sg2-BFP, sg3-BFP, and sg4-BFP also produced lower reporter signals than the non-targeting control. Cas13d-mediated BFP targeting showed a similar guide-dependent pattern, with sg1-BFP, sg2-BFP, and sg3-BFP producing the strongest reductions, while sg4-BFP showed comparatively weaker suppression. Together, these results demonstrate that reporter suppression depended on the Cas13 variant, guide sequence, and relative expression levels of the reporter and Cas13 proteins ^29^ **(Figure 3b)**.

To further compare Cas13b and Cas13d across additional reporter systems, we evaluated Renilla luciferase and firefly luciferase (Fluc) reporters **(Figure 3c)**. For Renilla targeting, all four Cas13b-associated guides reduced luminescence relative to the corresponding non-targeting control, although the magnitude of suppression varied among guides. Cas13d-mediated targeting produced more pronounced reductions, with luminescence signals approaching background levels under both tested expression conditions. The Fluc reporter assays also revealed guide-dependent differences in targeting activity. For both Cas13b and Cas13d, the extent of Fluc suppression varied among guides and between the lower-reporter/higher-Cas13 and higher-reporter/lower-Cas13 conditions. These findings indicate that both Cas13 variants can mediate reporter suppression, although their activity depends on the reporter target, guide sequence, and relative expression levels.

Having established reporter-targeting activity, we next evaluated whether Cas13d could target endogenous host transcripts. Four guide RNAs were assessed for each of five targets: human CTLA-4 (hCTLA-4), human PD-L1 (hPD-L1), mouse PD-L1 (mPD-L1), human TAZ (hTAZ), and human YAP (hYAP). Western blot analysis revealed guide-dependent differences in the abundance of the corresponding target proteins compared with the non-targeting controls (Fig. 3d). For hCTLA-4, individual guides produced variable reductions in protein abundance. Reduced PD-L1 protein signals were also observed following targeting of both human and mouse PD-L1. Similarly, targeting of TAZ and YAP resulted in reduced protein signals compared with the corresponding non-targeting controls. β-Actin was used as a loading control, and Ponceau S staining was included to assess total protein loading and membrane transfer. These results demonstrate the applicability of the Cas13d system to targeting multiple endogenous host transcripts, including genes encoding immune-regulatory and signaling-associated proteins.

We subsequently optimized the promoter used to drive guide RNA expression. Five RNA polymerase III promoters, U6, p5S, pOSU3, pH1, and p7SK, were compared using a BFP reporter-based assay **(Figure 3e)**. Reporter fluorescence was examined under both lower and higher BFP expression conditions. Differences in fluorescence intensity were observed among the promoter configurations, indicating that promoter selection influenced the functional activity of the guide RNA expression system. Among the promoters tested, p7SK showed strong reporter suppression and was selected for subsequent incorporation into the engineered HSV constructs.

Finally, the selected guide RNA expression configuration was evaluated in an HSV-based construct. HEK293 cells were examined following expression of HSV-U3-BFP/U37-Cas13d constructs containing either an ICP34.5-7SK-scrRNA cassette or an ICP34.5-7SK-sgBFP cassette. Fluorescence microscopy revealed a strong BFP signal in the non-targeting control condition, whereas substantially reduced fluorescence was observed in cells expressing the BFP-targeting guide **(Figure 3f)**. This result provided a functional demonstration of guide-dependent reporter suppression in the context of the engineered HSV construct.

These experiments demonstrated guide-dependent RNA-targeting activity of the Cas13 platform across multiple reporter systems and showed reduced protein abundance for several endogenous targets following Cas13d targeting. The promoter-screening experiments identified p7SK as the selected promoter for guide RNA expression, and the HSV-based construct further demonstrated BFP reporter suppression by the targeting guide. These findings provided the basis for incorporating the optimized Cas13d and p7SK guide RNA components into the oncolytic HSV platform for subsequent evaluation in tumor cell models ^25^.

### Establishment and validation of doxycycline-inducible expression systems in HEK293 cells and recombinant HSV

Following the functional characterization of Cas13d and optimization of the gRNA expression system, we next established a doxycycline-responsive regulatory platform to enable temporal control of transgene expression in the HSV background. Two complementary expression configurations based on the tetracycline-responsive Tet-OFF and Tet-ON systems were developed. The Tet-OFF constructs employed the tetracycline-controlled transactivator (tTA), whereas the Tet-ON constructs utilized the reverse tetracycline-controlled transactivator (rtTA3). For each regulatory configuration, three construct architectures were generated, containing zero, one, or two copies of the US11 gene. All constructs incorporated a Tet operator-controlled luciferase-P2A-GFP reporter cassette and an IRFP-720-P2A-NeoR/KanR cassette for fluorescent detection and antibiotic selection **(Figure 4a)**. The regulatory mechanisms of the two systems are illustrated in (**Figure 4b**). In the Tet-OFF system, transcription is activated in the absence of doxycycline and repressed upon doxycycline addition. Conversely, the Tet-ON system requires doxycycline for transcriptional activation ^31,38^. These configurations provided a framework for evaluating doxycycline-dependent reporter expression across different construct architectures.

**Figure 4.**
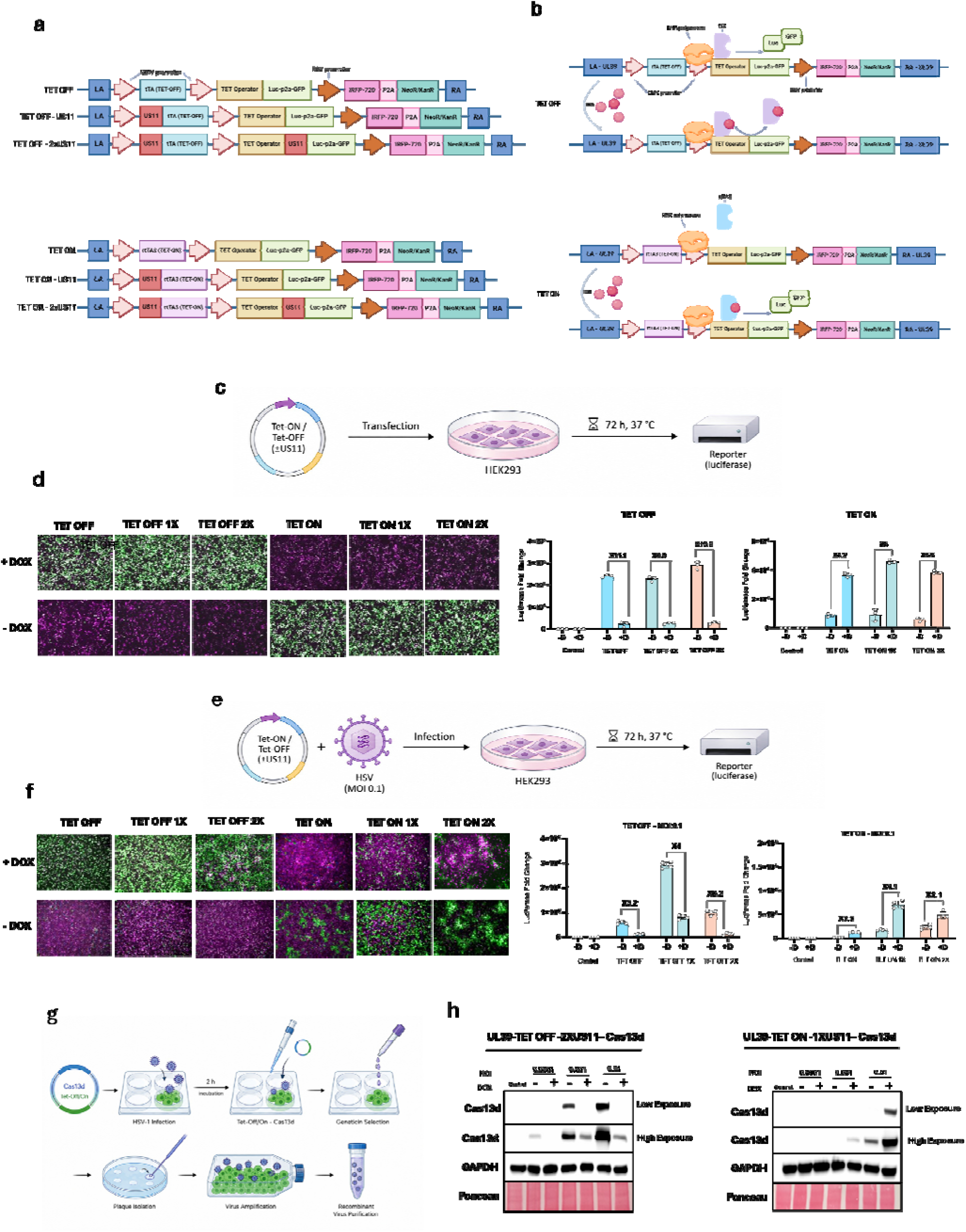
Development and validation of doxycycline-inducible expression systems in HEK293 cells and recombinant HSV. (a) Schematic representation of Tet-OFF and Tet-ON constructs containing zero, one, or two copies of US11. (b) Schematic illustration of doxycycline-dependent regulation in Tet-OFF and Tet-ON systems. (c) Experimental workflow for evaluating doxycycline-inducible reporter expression following transfection of HEK293 cells. (d) Representative fluorescence images and luciferase measurements under doxycycline-treated (+DOX) and untreated (−DOX) conditions following 72 h incubation. (e) Experimental workflow for evaluating doxycycline-dependent reporter expression following recombinant HSV infection at an MOI of 0.1. (f) Representative fluorescence images and luciferase measurements following 72 h incubation at 37 °C under +DOX and −DOX conditions. (g) Workflow for recombinant HSV-Cas13d generation, selection, amplification, and purification. (h) Western blot analysis of Cas13d expression in representative Tet-OFF and Tet-ON recombinant HSV constructs at MOIs of 0.0001, 0.001, and 0.01, under +DOX and −DOX conditions. GAPDH and Ponceau S were used as loading and transfer controls, respectively.

We first evaluated the regulatory activity of the constructs in HEK293 cells following plasmid transfection **(Figure 4c)**. Cells were transfected with the corresponding Tet-OFF or Tet-ON constructs containing zero, one, or two copies of US11 and incubated for 72 h at 37 °C in the presence or absence of doxycycline (DOX). Reporter expression was assessed by fluorescence microscopy and luciferase assays **(Figure 4d)**.

The Tet-OFF configurations exhibited the expected inverse response to doxycycline, with substantially higher reporter activity in the absence of DOX and reduced activity following DOX treatment. The differences in reporter activity between the two conditions were approximately 11.1-fold, 8.9-fold, and 10.8-fold for the Tet-OFF, Tet-OFF-1×US11, and Tet-OFF-2×US11 constructs, respectively. This regulatory response was maintained across all three architectures, indicating that the presence of additional US11 copies did not abolish doxycycline-dependent repression.

The Tet-ON configurations showed the opposite response, with relatively low reporter activity in the absence of DOX and increased activity following doxycycline treatment. The corresponding increases were approximately 4.2-fold, 5.0-fold, and 6.6-fold for the Tet-ON, Tet-ON-1×US11, and Tet-ON-2×US11 constructs, respectively (Fig. 4d). Together, these results confirmed the functionality of both inducible regulatory systems in HEK293 cells and demonstrated that doxycycline-dependent control was maintained across the different US11 configurations.

Having established inducible reporter regulation following transfection, we next investigated whether these regulatory systems retained their activity in the context of HSV infection. HEK293 cells were infected with the corresponding recombinant HSV constructs at an MOI of 0.1 and incubated for 72 h at 37 °C in the presence or absence of DOX **(Figure 4e)**. Reporter expression was subsequently evaluated by fluorescence microscopy and luciferase assays **(Figure 4f)**.

The Tet-OFF constructs maintained their characteristic doxycycline-dependent repression profile during HSV infection, with higher reporter activity in the absence of DOX than in its presence. The differences in reporter activity were approximately 5.2-fold, 4.0-fold, and 6.2-fold for the Tet-OFF, Tet-OFF-1×US11, and Tet-OFF-2×US11 constructs, respectively. Among these configurations, Tet-OFF-2×US11 exhibited the largest differential response.

Similarly, the Tet-ON constructs retained doxycycline-dependent activation in the viral context. Reporter activity increased following DOX treatment, with approximately 7.5-fold, 4.1-fold, and 2.1-fold increases observed for the Tet-ON, Tet-ON-1×US11, and Tet-ON-2×US11 constructs, respectively **(Figure 4f)**. Although the magnitude of induction differed among the constructs and compared with the transfection-based experiments, both systems maintained their expected doxycycline-dependent regulatory behavior during HSV infection. These findings demonstrated that the inducible regulatory elements remained functional within the viral context and were not restricted to plasmid-based expression ^39^.

Having established doxycycline-responsive regulation in both transfection-based and virus-mediated expression systems, we next generated recombinant HSV constructs encoding Cas13d. As illustrated in Fig. 4g, the production workflow involved infection with parental HSV-1, followed by a 2 h incubation and subsequent transfection with the corresponding Tet-OFF-or Tet-ON-regulated Cas13d construct. Transfected cultures were subjected to Geneticin selection, followed by plaque isolation, viral amplification, and purification of the resulting recombinant viruses ^40^.

To further assess doxycycline-dependent Cas13d expression at the protein level, two representative recombinant constructs, UL39-Tet-OFF-2×US11-Cas13d and UL39-Tet-ON-1×US11-Cas13d, were evaluated at three different MOIs (0.0001, 0.001, and 0.01) in the presence or absence of DOX **(Figure 4h)**. Western blot analysis revealed that Cas13d protein abundance varied with viral input and doxycycline treatment. The Tet-OFF construct showed preferential Cas13d expression in the absence of DOX, whereas the Tet-ON construct exhibited stronger Cas13d signals under doxycycline-treated conditions. Both low-and high-exposure images are presented to visualize Cas13d expression across the different infection conditions. GAPDH was used as a loading control, and Ponceau S staining was included to assess total protein loading and membrane transfer.

The comparison of the two regulatory configurations established that doxycycline-dependent control was retained after transfer into the viral context. Notably, Tet-OFF-2×US11 showed the largest differential response among the Tet-OFF configurations during HSV infection and was therefore selected for subsequent control of Cas13d expression. This selected configuration provided the inducible viral platform for evaluating Cas13d-mediated modulation of host antiviral responses in tumor cell models.

### Screening of target-specific HSV-iCas13d variants reveals cell-dependent fluorescence patterns

Following the establishment of the inducible HSV-iCas13d platform, we next investigated the effects of targeting selected host antiviral and signaling-associated factors on fluorescence patterns during HSV infection. Based on the interconnected nature of innate immune and antiviral pathways, we designed guide RNAs targeting 15 host factors: TBK1, TRAF6, TRAF3, JAK1, STAT1, PKR, IRF3, IRF7, MYD88, JNK, IKKβ, IFNB, RIG-I, MAVS, and NEDD8 **(Figure 5a)** ^36^. These targets represent different components of cellular signaling networks involved in innate immune recognition, interferon responses, inflammatory signaling, and cellular stress regulation.

**Figure 5.**
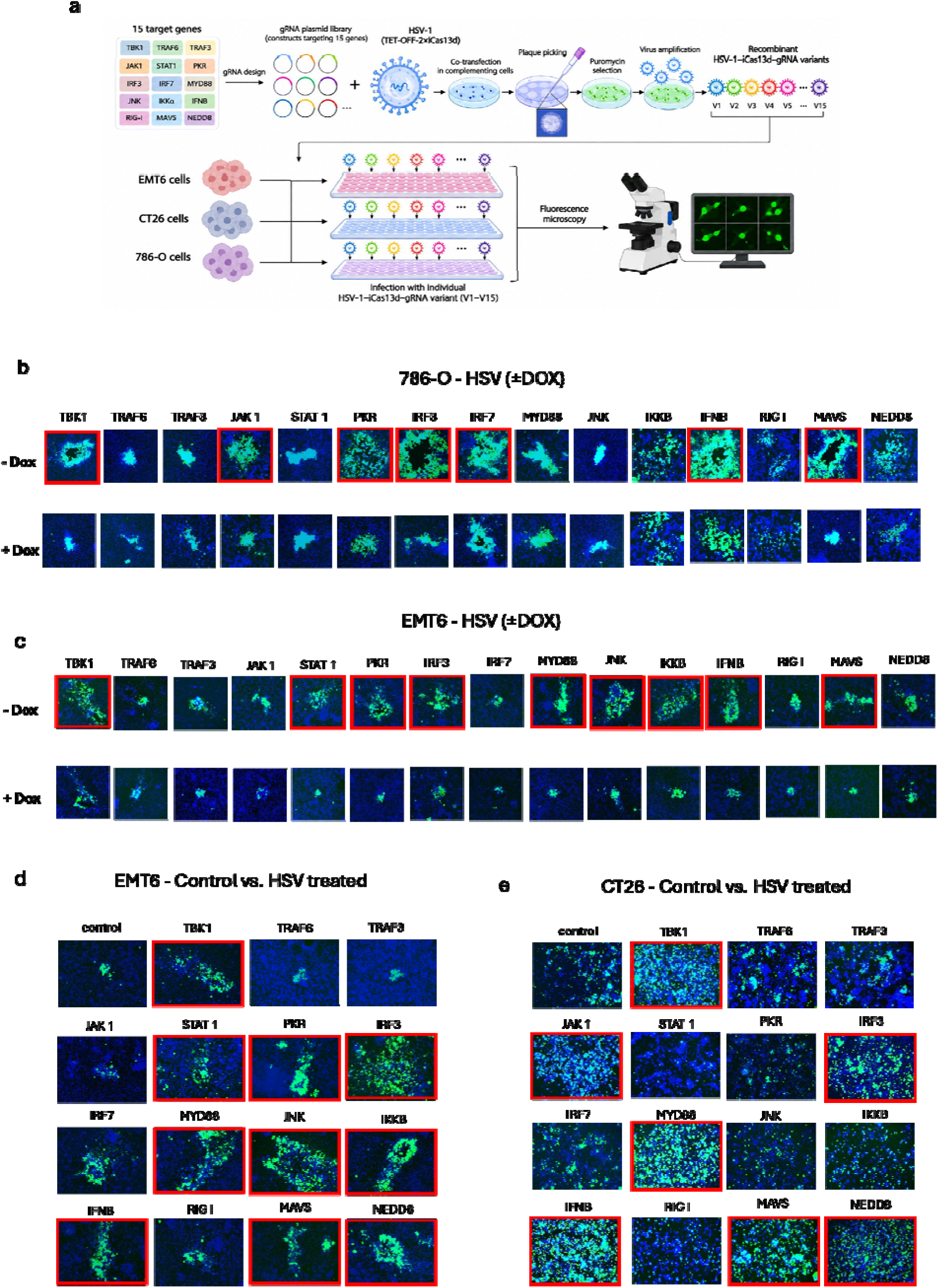
Generation and screening of target-specific HSV-iCas13d variants across tumor cell models. (a) Schematic overview of gRNA design targeting 15 host antiviral and signaling-associated targets and generation of recombinant HSV-iCas13d-gRNA variants (V1–V15). (b,c) Representative fluorescence images of 786-O and EMT6 cells, respectively, following HSV infection under doxycycline-treated (+DOX) and untreated (−DOX) conditions. (d,e) Representative fluorescence images comparing control and HSV-treated EMT6 and CT26 cells, respectively. Red boxes highlight selected target-specific conditions with visually distinct fluorescence patterns. Images were acquired 72 h post-infection under identical imaging conditions. Fluorescence patterns were evaluated qualitatively.

The corresponding gRNA constructs were assembled into a plasmid library and introduced together with the engineered HSV-1 backbone encoding Tet-OFF-regulated iCas13d. Recombinant viral variants were generated through co-transfection in complementing cells, followed by plaque picking, puromycin selection, and viral amplification. This workflow resulted in a collection of target-specific HSV-1-iCas13d-gRNA variants, which were subsequently evaluated in three tumor cell models: murine EMT6 and CT26 cells and human 786-O cells **(Figure 5a)**. Individual viral variants were used to infect each cell model, and reporter-associated fluorescence was examined by fluorescence microscopy.

We initially examined the fluorescence patterns associated with the individual targeting variants in 786-O and EMT6 cells under doxycycline-treated and untreated conditions **(Figure 5b,c)**. In both cell models, differences in fluorescence intensity and distribution were observed among the individual targeting conditions. However, the patterns varied between the two cellular backgrounds, suggesting that the fluorescence-associated responses to individual gRNA constructs were dependent on the cell model.

In 786-O cells, selected fluorescence patterns were highlighted by red outlines **(Figure 5b)**. These included the conditions corresponding to TBK1, JAK1, PKR, IRF3, IRF7, IFNB, and MAVS. In EMT6 cells, the highlighted conditions included TBK1, STAT1, PKR, IRF3, MYD88, JNK, IKKβ, IFNB, and MAVS **(Figure 5c)**. The differences in the highlighted conditions between the two cell lines indicate that the fluorescence patterns associated with individual targeting constructs were not uniform across the tested tumor models.

Among the selected host factors, JNK-associated signaling represents an interesting candidate for further investigation. Unlike several classical antiviral signaling components, JNK belongs to the stress-activated mitogen-activated protein kinase (MAPK) family and has also been implicated in supporting HSV-1 lytic infection. Previous studies have shown that HSV-1 can activate JNK signaling and that this pathway may contribute to viral gene expression and replication ^41^. In our screening, the JNK-targeting condition was highlighted in EMT6 cells in both HSV-related comparisons, whereas a similar highlighted pattern was not observed in the 786-O or CT26 cells. Although these fluorescence observations do not establish a direct functional relationship between JNK targeting and viral replication, they identify JNK-associated signaling as a candidate for further investigation in the EMT6 model.

Additional fluorescence comparisons involving EMT6 and CT26 cells are presented in the lower image panels **(Figure 5d,e)**. Red outlines identify selected fluorescence patterns in these comparisons. The distribution of highlighted conditions differed between the two cell models, further illustrating the variability of fluorescence-associated responses across tumor cell backgrounds. These observations support the importance of evaluating target-specific HSV-iCas13d variants in multiple cellular models rather than relying on a single cell line.

These experiments established a fluorescence-based screening framework for evaluating target-specific HSV-iCas13d variants across EMT6, CT26, and 786-O tumor cell models. The distinct fluorescence patterns observed among individual targeting conditions and cellular backgrounds highlight the complexity of host-virus interactions and indicate that the responses associated with individual host targets may be cell-dependent. The red-outlined conditions were selected based on their observed fluorescence patterns and provide candidates for subsequent investigation. However, fluorescence microscopy alone does not establish successful suppression of the corresponding host targets or changes in infectious virus production. Further assessment of target transcript and protein abundance, together with infectious viral titration, will therefore be necessary to determine the functional consequences of individual gRNA constructs.

## CONCLUSION

This study presents the development of an inducible, HSV-encoded CRISPR-Cas13d platform for programmable modulation of virus–host interactions. By integrating transcriptomic profiling and pharmacological screening, we identified cell-dependent differences in host antiviral responses and demonstrated that pharmacological inhibition of selected pathways could enhance infectious HSV production across multiple tumor cell models. We further characterized Cas13-mediated RNA targeting, optimized gRNA expression using the p7SK promoter, and established doxycycline-responsive Tet-ON and Tet-OFF systems to achieve controlled Cas13d expression within the HSV context. The generation and initial screening of HSV-iCas13d variants targeting 15 host antiviral and signaling-associated factors revealed distinct fluorescence patterns across EMT6, CT26, and 786-O cells, highlighting the influence of cellular context on host–virus interactions. Although further molecular and functional validation is required to establish the effects of individual gRNAs on target suppression and viral replication, this platform provides a versatile foundation for investigating host antiviral determinants and developing more selective strategies to modulate oncolytic HSV activity. Ultimately, this approach may contribute to the development of programmable oncolytic virus systems that balance viral replication with host immune responses.

## Methods

### Cell Culture

Human, murine, and African green monkey cell lines, including U2OS, Vero, HEK293, 786-O, EMT6, CT26, 67NR, and AT3, were used in this study. All cell lines were obtained from the American Type Culture Collection (ATCC). Cells were cultured in Dulbecco’s Modified Eagle Medium (DMEM) supplemented with 10% heat-inactivated fetal bovine serum (FBS) and 1% penicillin–streptomycin. Cell cultures were maintained at 37 °C in a humidified atmosphere containing 5% CO. Cells were routinely passaged using trypsin–EDTA and washed with Dulbecco’s phosphate-buffered saline (DPBS), as required. Cell cultures were regularly monitored for mycoplasma contamination using an external testing service.

### Viruses

An HSV-based platform was engineered to express the CRISPR-Cas13d system under doxycycline-responsive regulation. The recombinant HSV constructs contained a doxycycline-regulated Cas13d expression cassette, a guide RNA (gRNA) expression cassette, and a luciferase-P2A-GFP reporter cassette.

The Cas13d expression cassette was inserted into the UL39 locus of the HSV genome under the control of either Tet-OFF or Tet-ON regulatory elements. Both configurations were generated and evaluated to compare doxycycline-dependent Cas13d expression. Different viral configurations containing zero, one, or two copies of the US11 gene were also evaluated during construct optimization. Based on these comparative analyses, the Tet-OFF configuration containing two copies of US11 (Tet-OFF-2×US11; OFF2x) was selected for subsequent experiments. The gRNA expression cassette was inserted into the ICP34.5 locus under the control of the human 7SK promoter.

The engineered HSV constructs shared the same general genomic architecture and differed primarily in their gRNA sequences. The gRNA library included guides targeting 15 host antiviral and signaling-associated targets, together with two non-targeting control sequences.

### Construct Design and gRNA Selection

Plasmid and viral genome designs were generated using SnapGene software. Guide RNAs targeting 15 host antiviral and signaling-associated targets were designed using the CRISPR-Cas13 design platform developed by the New York Genome Center (https://cas13design.nygenome.org/), followed by manual evaluation and selection. Candidate gRNAs were assessed based on the design scores and predicted off-target information provided by the platform. Guides were designed against the corresponding human and mouse target sequences. Two distinct non-targeting gRNAs were included as negative controls.

### Reagents and Small-Molecule Inhibitors

Doxycycline hyclate (Sigma-Aldrich, catalog no. D9891-25G) was used to regulate the inducible expression system at a final concentration of 1 µg/mL. Puromycin (Thermo Fisher Scientific, catalog no. A1113803) and Geneticin (G418; Thermo Fisher Scientific, catalog no. 10-131-035) were used for antibiotic selection during the generation of engineered constructs. Puromycin and Geneticin were used at final concentrations of 1 µg/mL and 500 µg/mL, respectively.

Thirteen small-molecule inhibitors associated with host antiviral signaling pathways were evaluated in this study: MRT67307, GSK8612, momelotinib, C25-140, amlexanox, PKR inhibitor (PKRi), BX795, dimethyl fumarate, pevonedistat, BAY 11-7082, 4-octyl itaconate (4-OI), TPCA-1, and ruxolitinib. These compounds were selected to investigate the contribution of different host signaling pathways to the regulation of HSV infection.

All inhibitors were evaluated at a final concentration of 0.1 µM. Dimethyl sulfoxide (DMSO) was used as the vehicle control.

### Virus Production and Titration

Vero cells were used for the production of HSV stocks. Cells were infected at a multiplicity of infection (MOI) of 0.1 and maintained in roller bottles containing 250 mL of culture medium supplemented with 10% FBS. Infected cultures were maintained for 72 h, after which both cells and culture supernatants were harvested. The collected material was subjected to five freeze–thaw cycles to facilitate viral release.

Following freeze–thaw treatment, the culture supernatant and cell-associated material were processed for viral recovery. The culture supernatant was subjected to ultracentrifugation through a 36% sucrose cushion using an Optima XE ultracentrifuge equipped with an SW32Ti rotor and Beckman ultracentrifuge tubes (catalog no. 344058). Centrifugation was performed at 25,000 rpm for 2 h at 4 °C. The resulting viral pellet was combined with the cell-associated pellet and subjected to a second centrifugation step under the same conditions for an additional 2 h. The recovered viral pellet was resuspended in sterile phosphate-buffered saline (PBS) and stored at −80 °C until use.

Viral titers were determined by plaque assay using U2OS cells. A total of 1.2 × 10 U2OS cells were seeded in 24-well plates (5 × 10 cells per well). Serial tenfold dilutions of viral stocks were prepared and added to the cell monolayers. Following a 2-h adsorption period, the inoculum was replaced with an overlay consisting of equal volumes of 3% carboxymethyl cellulose (CMC) and 2× DMEM supplemented with FBS. Plates were incubated for 72 h at 37 °C in a humidified atmosphere containing 5% CO. Viral plaques were visualized by crystal violet staining for 3 min. Viral titers were calculated and expressed as plaque-forming units per milliliter (PFU/mL). Each viral titration was performed in triplicate.

### Small-Molecule Inhibitor Treatment and Viral Infection

To evaluate the effects of pharmacological inhibition of host antiviral pathways on HSV infection, AT3, 67NR, EMT6, CT26, and 786-O cells were pretreated with the indicated small-molecule inhibitors, using DMSO-treated cells as vehicle controls. Following pretreatment, cells were infected with HSV at an MOI of 0.1 and maintained for 72 h post-infection. Viral reporter fluorescence was evaluated by fluorescence microscopy, and infectious viral production was quantified by plaque assay. Viral titers were expressed as plaque-forming units per milliliter (PFU/mL). Experiments were performed using three independent biological replicates.

### Western Blot Analysis

Protein expression was evaluated by Western blotting. Briefly, cells were lysed using Cell Lysis Buffer II, and total protein concentrations were determined using a bicinchoninic acid (BCA) protein assay. Equal amounts of protein were separated by SDS-PAGE and transferred onto nitrocellulose membranes. Membranes were stained with Ponceau S to assess total protein loading and transfer efficiency, followed by blocking with 5% skim milk.

The membranes were incubated with primary antibodies against CTLA-4, PD-L1, TAZ, YAP, Cas13d, β-actin, and GAPDH, as appropriate for each experiment. Following incubation with the corresponding secondary antibodies, protein bands were detected using an enhanced chemiluminescence (ECL) detection system. β-Actin and GAPDH were used as loading controls.

### Fluorescence Microscopy

Fluorescence imaging was performed using an EVOS M5000 imaging system. Images were acquired at the indicated experimental endpoints to evaluate reporter expression and infection-associated fluorescence. For infection-based experiments, images were acquired 72 h post-infection. GFP and BFP signals were examined at 4× and 10× magnifications. Exposure settings were kept identical across experimental groups to allow qualitative comparison of fluorescence patterns.

Three images were acquired per well, and experiments were performed in three replicates. Fluorescence signals were assessed qualitatively, and no quantitative fluorescence intensity analysis was performed.

### Comparison of Cas13b and Cas13d Activity

The functional activities of Cas13b and Cas13d were compared in HEK293 cells using reporter-based assays. Cells were transfected with plasmids encoding either Cas13b or Cas13d and the corresponding reporter constructs. The reporter systems included blue fluorescent protein (BFP), NanoLuc, firefly luciferase, and Renilla luciferase.

Reporter activity was evaluated using fluorescence-based and luminescence-based readouts, as appropriate for each reporter system. The results were used to compare the functional performance of the two Cas13 variants. Experiments were performed in three replicates.

### Optimization of gRNA Expression Promoters

To identify an effective promoter for gRNA expression, five promoters were evaluated: U6, p5S, pOSU3, pH1, and p7SK. The promoter constructs were designed using the same general backbone and the same gRNA sequence to allow comparison of promoter performance while minimizing sequence-dependent differences.

Promoter activity was evaluated using the corresponding reporter-based functional readout. Based on the comparative results, the p7SK promoter was selected for subsequent engineering of the HSV constructs.

### Evaluation of Doxycycline-Regulated Expression Systems

Tet-ON and Tet-OFF configurations were evaluated in HEK293 cells to establish doxycycline-dependent regulation of the engineered expression system. Both configurations, containing zero, one, or two copies of US11, were evaluated in the presence or absence of doxycycline at a final concentration of 1 µg/mL. Reporter expression was assessed following plasmid transfection and recombinant HSV infection. For infection-based experiments, cells were infected at an MOI of 0.1 and incubated for 72 h.

System performance was evaluated using fluorescence microscopy, luciferase measurements, and Western blot analysis. The regulatory responses of the Tet-ON and Tet-OFF configurations were compared under doxycycline-treated and untreated conditions. Based on these comparative evaluations, the Tet-OFF-2×US11 configuration was selected for subsequent regulation of Cas13d expression in the engineered HSV constructs.

### Evaluation of Engineered HSV Constructs in Tumor Cell Models

The engineered HSV-iCas13d constructs were evaluated in the murine tumor cell lines EMT6 and CT26 and the human tumor cell line 786-O. Cells were infected with individual gRNA-containing HSV constructs at the same MOI across experimental groups and maintained in the presence or absence of doxycycline, as indicated. Reporter-associated fluorescence was examined 72 h post-infection by fluorescence microscopy to compare the patterns associated with individual targeting constructs across the different tumor cell models. Each targeting construct was evaluated alongside the corresponding non-targeting control to identify cell-dependent differences in fluorescence patterns.

### RNA Sequencing and Bioinformatic Analysis

RNA sequencing was performed to characterize host transcriptional responses to oHSV-1 infection in five cancer cell lines, including four murine cancer cell lines (EMT6, CT26, AT3, and 67NR) and one human cancer cell line (786-O). The primary transcriptomic analysis presented in Figure 1 focused on the four murine cell lines, whereas data from 786-O were included in the supplementary apoptosis-related pathway analysis. For each cell line, two experimental conditions were evaluated: uninfected control and HSV-infected cells. Cells in the infection groups were exposed to HSV-1 at a multiplicity of infection (MOI) of 1 and incubated for 72 h at 37 °C under 5% CO. Following incubation, cells were harvested and stabilized using DNA/RNA Shield (Zymo Research). Samples were subsequently submitted to Plasmidsaurus for RNA extraction, library preparation, sequencing, and initial bioinformatic processing.

RNA sequencing was performed using a 3′ end-counting approach based on poly(A) mRNA capture. Transcripts were captured using oligo-dT primers and converted into cDNA through reverse transcription and second-strand synthesis. Libraries were prepared by tagmentation, amplified using unique dual indices, and sequenced on an Illumina NovaSeq instrument, generating approximately 90-bp reads. Unique molecular identifiers (UMIs) were used to remove PCR and optical duplicates.

Filtered reads were aligned to the corresponding mouse or human reference genome using STAR and coordinate-sorted using samtools v1.21. Gene-level expression was quantified using the provider’s standardized count-based workflow. Normalized expression values and differential gene expression results were generated by comparing HSV-infected samples with their corresponding uninfected controls for each cell line.

For pathway analysis, genes were ranked according to their log fold changes and analyzed using preranked Gene Set Enrichment Analysis (GSEA) in Python with GSEApy v1.1.10 and the species-specific Reactome 2022 collections (Mouse or Human). Gene sets containing 5–5000 genes were included, using 1,000 permutations and a fixed random seed of 42. Normalized enrichment scores (NES) and false discovery rate (FDR)-adjusted P values were extracted.

Heatmaps were generated from log-transformed normalized expression values and converted into row-wise Z-scores. Genes associated with innate immune responses, interferon signaling, cytokine signaling, antiviral responses, and other pathways identified through GSEA were selected for visualization. For the combined murine heatmap, genes belonging to the Mouse Reactome 2022 Immune System pathway were retrieved using GSEApy and intersected with the selected gene set. Enrichment plots, heatmaps, and summary figures were generated using GSEApy v1.1.10, Python, and GraphPad Prism 11.0.2.

## ABBREVIATIONS

HSV: Herpes Simplex Virus
HSV-1: Herpes Simplex Virus Type 1
oHSV: Oncolytic Herpes Simplex Virus
CRISPR: Clustered Regularly Interspaced Short Palindromic Repeats
Cas13d: CRISPR-associated protein 13d
gRNA: Guide RNA
MOI: Multiplicity of Infection
DOX: Doxycycline
GFP: Green Fluorescent Protein
BFP: Blue Fluorescent Protein
PFU: Plaque-Forming Unit
RNA-seq: RNA Sequencing
GSEA: Gene Set Enrichment Analysis
NES: Normalized Enrichment Score
FDR: False Discovery Rate.

## AUTHOR CONTRIBUTIONS

Conceptualization: Zahra Ameli, Reza Rezaei, Taha Azad; Methodology: Zahra Ameli, Reza Rezaei, Taha Azad; Investigation: Zahra Ameli, Reza Rezaei, Taha Azad; Formal analysis: Zahra Ameli, Zumama Khalid, Reza Rezaei, Taha Azad; Data curation: Zahra Ameli, Reza Rezaei; Writing – original draft: Zahra Ameli; Writing – review and editing: All authors; Supervision: Taha Azad; Funding acquisition: Taha Azad. Supported by start-up funding from Université de Sherbrooke. All authors have read and approved the final manuscript.

## ACKNOWLEDGMENT

The authors would like to thank the members of the Azad laboratory and the research facilities at Université de Sherbrooke for their technical support and assistance throughout this study. The authors also gratefully acknowledge the Université de Sherbrooke Cancer Research Institute (IRCUS) for financial support. Finally, the authors thank Abril Ixchel Munoz Zuniga for her help.

